# MS/MS Mass Spectrometry Filtering Tree for Bile Acid Isomer Annotation

**DOI:** 10.1101/2025.03.04.641505

**Authors:** Ipsita Mohanty, Shipei Xing, Vanessa Castillo, Julius Agongo, Abubaker Patan, Yasin El Abiead, Helena Mannochio-Russo, Simone Zuffa, Jasmine Zemlin, Alexandre Tronel, Audrey Le Gouellec, Thomas Soranzo, Crystal X. Wang, Jennifer E. Iudicello, Mohammadsobhan S. Andalibi, Ronald J. Ellis, David J. Moore, Donald R. Franklin, Marta Sala-Climent, Monica Guma, Robin M. Voigt, Rima Kaddaruh-Dauok, Dionicio Siegel, Mingxun Wang, Lee R. Hagey, Pieter C. Dorrestein

## Abstract

Bile acids are essential steroids regulating immunity, nutrient absorption, insulin, appetite, and body temperature. Their structural diversity is vast, but due to spectral similarities, MS/MS spectral matching often fails to resolve isomers. This study introduces a proof-of-concept workflow using a mass spectrometry query language filtering tree that distinguishes isomeric bile acids in untargeted LC-MS/MS data. Its application revealed a deoxycholyl-2-aminophenol amidate linked to whole grain consumption.

## Main text

Cholesterol-derived steroid molecules, known as bile acids, have been detected in nearly every organ where they have been studied^1^. Alterations in bile acid composition and dysbiosis have been linked to various health conditions, including neurocognitive developmental disorders, cancer, infections, and metabolic diseases such as diabetes and IBD^2^. There is a growing recognition that the structural diversity of bile acids has been vastly underestimated^1,3–19^. Despite the recent structural annotation of thousands of bile acids, compiling an inventory of all bile acids and their modified forms remains a considerable challenge, underscoring the need for innovative methods to capture this diversity. One of the main challenges is the vast diversity of theoretically possible bile acid isomers.

In humans, bile acids are predominantly derived from cholesterol through pathways that shorten the side chain, producing a carboxyl terminus on the remaining 24-carbon steroid core, which consists of four interconnected rings, collectively referred to here as the “core” (colored in **Figure 1a**). The core undergoes hydroxylation, a process typically associated with the liver, but it can also occur in other organs such as the brain, kidney, and spleen^5^. This process primarily produces cholic and chenodeoxycholic acids. Once exposed to the microbiome, the steroid core of host-derived bile acids undergoes multiple chemical modifications catalyzed by microbial enzymes, including hydroxylation, dehydroxylation, dehydration, and oxidation of hydroxyl functional groups^17,20–22^. Considering hydroxylation at known carbon positions on bile acids reported from human samples - C1, C3, C4, C6, C7, C11, C12, C15, C16, C22, and C23 (as numbered in **Figure 1a, Supplementary Table S1)** - along with stereoisomers and allo versus non-allo A/B ring isomerization at C5, approximately 3,124 candidate mono-, di-, and tri-hydroxy bile acid cores are theoretically possible (**Figure 1a**). If additional hydroxylation positions are identified in the future, the total count will increase accordingly. The carboxylate group on the bile acid side chain can undergo amidation, and with the expanding variety of possible amidations for each core structure^1^, the total number of core-conjugate combinations likely extends to hundreds of thousands, possibly even millions, especially when the conjugations on hydroxylations are added to this equation^5^. While progress is being made in expanding carboxylate modifications, characterizing the bile acid core structure remains a key challenge in LC-MS/MS-based untargeted metabolomics. Alternative and innovative strategies using ion mobility mass spectrometry^23–25^, statistical probability^26^ and MS/MS fragmentation^27,28^ have been developed; however, neither approach is yet compatible with untargeted LC-MS/MS datasets and re-analysis of public data.

**Figure 1.**
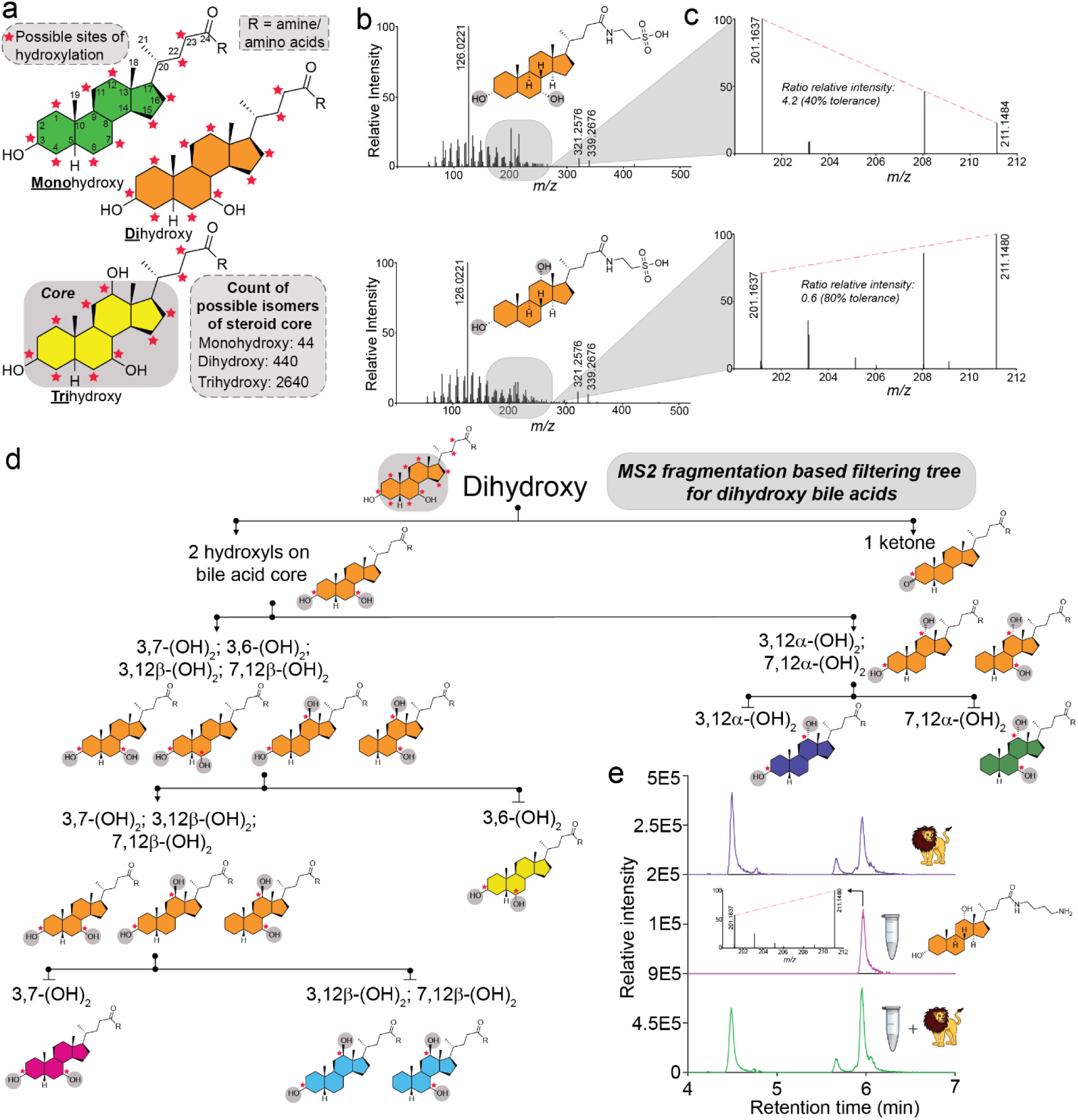
Development of MS/MS Fragmentation-Based MassQL Filters. **a)** Structure of bile acids highlighting mono-, di-, and tri-hydroxylated steroid cores, with experimentally observed potential hydroxylation sites on the steroid core indicated by red stars. **b)** MS/MS fragmentation spectra of the regioisomers, taurochenodeoxycholic acid and taurodeoxycholic acid, illustrating a low-intensity mass region containing ions unique to each isomer. **c)** Enlarged view of the MS/MS fragmentation spectra for taurochenodeoxycholic and taurodeoxycholic acids, emphasizing the ion pair used to calculate relative intensity ratios for differentiating these isomers. **d)** Sequential MS/MS fragmentation-based filtering tree designed to classify regio- and stereoisomers of dihydroxylated bile acids. Structures at each filtering step are shown, with terminal bins color-coded for clarity. **e)** Retention time and MS/MS analysis confirmed the MassQL tree-predicted core structure of the bile acid conjugate, deoxycholyl-putrescine by comparing with a synthetic standard.

Untargeted LC-MS/MS analyses typically rely on MS/MS spectral reference libraries to identify bile acids. However, MS/MS spectral alignment with libraries generally cannot distinguish between bile acid isomers, as their stable four-ring core structure leads to similar fragmentation patterns and MS/MS fragment ions. Recently, we developed MS/MS fragmentation-based filters using the MassQL data pattern filtering strategy^29^ to retrieve and create a reference library of ∼21,000 MS/MS spectra that represent detectable bile acid signatures to enhance the discovery of bile acids in untargeted metabolomics experiments, irrespective of the ion form^1^. This library includes MS/MS spectra of known and many yet-to-be structurally characterized bile acid candidates that have already been detected from public metabolomics data repositories. This includes MS/MS of multiple ion forms such as different adducts, in-source fragments, and multimers^1^. The MassQL library enables annotation of the number of hydroxyl groups on the bile acid steroid core and any modifications present. However, the bile acid MS/MS fragments used to build this reference library for MassQL-based filtering do not allow differentiation between positional and stereoisomers.

MS/MS spectral matching often overlooks finer details within fragment ions, as these minimally influence overall spectral match scores (**Figure 1b**). We hypothesized that information in these low intensity MS/MS fragments, including specific ion pairs, intensity ratios, and unique ions, could be used to distinguish the steroid core isomers. To evaluate this, we developed a MassQL-based filtering tree utilizing key marker ions and their intensity ratios within narrow *m/z* windows to propose regio- and stereoisomer assignments of bile acids solely from MS/MS data. In this study, we focus on bile acid amidates, structures found in humans and research animals.

To create the filtering tree, we curated ion intensity patterns by manually inspecting an MS/MS spectral library of 48 mono-, di-, and tri-hydroxylated bile acid taurine amidates, as well as ketone-conjugated bile acids also conjugated with taurine **(Supplementary Table S1,** *see MS/MS data acquisition of taurine-conjugated bile acids in Methods*). MS/MS fragment ions that contributed minimally to spectral similarity scoring yet exhibited variability among isomers were selected. For example, taurochenodeoxycholic acid (TCDCA) and its regioisomer taurodeoxycholic acid (TDCA) revealed diagnostic ion pairs (*m/z* 201.163 and *m/z* 211.147) within the *m/z* 150–300 range. The intensity ratio of these ions reliably differentiated the C12α hydroxyl group from other steroid cores, including its C12β epimer (**Figure 1c**). The proximity of these ions in *m/z* and their consistency across instruments and collision energies make them candidate markers, allowing for the formulation of a structural hypothesis with respect to the regio and stereochemistry of bile acid cores.

Using these insights, we developed MassQL queries incorporating diagnostic intensity ratios to classify bile acids. To enhance specificity and reduce false discoveries, individual queries were integrated into a branched filtering tree, progressively narrowing classifications from broad bile acid categories to specific isomers. Separate trees addressed mono-, di-, and trihydroxylated bile acids (**Figure 1d, Supplementary Figure S1**). The first branch in the tree separated ketone bile acids, leveraging diagnostic ions (e.g., *m/z* 161.132 in a dihydroxylated tree) to distinguish them from non-ketone-containing bile acids. Notably, we observed that a mono ketone bile acid generates the same bile acid core ion fragments as those produced by two hydroxyl groups in the MS/MS spectrum (**Supplementary Figure S2**). Consequently, the characteristic dihydroxylated bile acid ions at *m/z* 321.26 and 339.27 were also detected in monohydroxylated bile acids with a ketone on the steroid core. Even though the precursor *m/z value*s differ by 2 Da, the diagnostic ions will group bile acids containing one ketone and two hydroxyl groups into the same dihydroxy bin. Subsequent queries refined the regio- and stereochemical core assignments by analyzing the intensity ratios of closely spaced ions within the spectrum. Specificity and false discovery rates (FDRs) for each query were evaluated using reference MS/MS spectra of stereochemically defined bile acids (n=152 for monohydroxy; n=856 for dihydroxy; n=1524 for trihydroxy), ensuring that the MassQL filters reliably distinguished isomers within acceptable FDR parameters (**Supplementary Table S2**). The large number of MS/MS spectra for bile amidates recovered indicated that our MassQL queries (developed with taurine bile amidates) also captured the bile acid isomer core for other amine-conjugated bile acids.

The utility of this approach was demonstrated by applying the isomer filters to the amidated MS/MS spectra that we previously uncovered from public data mining^1^, wherein we merged all MS/MS spectra with a cosine score higher than 0.7. As MS/MS spectra of different bile acid isomer cores, but with the same number of hydroxylations, were merged during the previous study^1^, we revisited the MS/MS spectra before merging. By analyzing these individual unmerged spectra, the original set of merged spectra for dihydroxy bile acids increased for some bile amidates, such as for tyrosine conjugated to a trihydroxy bile acid core (**Supplementary Table S2**). To highlight a specific example, we reanalyzed lion feces data from one of our previous studies, where we discovered polyamine-conjugated bile acids^1,30^. In that study, we had matched the retention times of one of the lower-intensity chromatographic peaks with the synthetic standard of chenodeoxycholyl-putrescine. However, the LC-MS trace also contained other isomers of bile acid-putrescine conjugates that were even more abundant. Using our MassQL filtering tree, we could predict that one of the more intense peaks in the LC-MS data was the deoxycholyl-putrescine amidate. This prediction of deoxycholyl-putrescine was validated by synthesizing the standard and matching the retention time (**Figure 1e**). In this example, the filtering tree helped to narrow down the candidates for steroidal core, thereby limiting the number of bile acids to be synthesized to confirm the structure.

To further illustrate the utility of the MS/MS filtering tree for bile acid isomer assignment, we re-analyzed a publicly available untargeted LC-MS/MS dataset comprising paired small intestinal fluid and feces samples from 15 healthy participants in France^31^. To get our first insight into the diversity of bile acids, we matched the MS/MS spectra against the GNPS spectral libraries, including the candidate bile acid MS/MS reference library that we had previously created from the repository scale analysis^1^. This revealed 1,499 MS/MS spectral matches to bile acids. We applied an additional filtering step to exclude MS/MS spectral matches to candidate bile acids where the experimental spectra lacked the two diagnostic MS/MS ions for bile acids^4^. Steroids and lipids often exhibit similar MS/MS spectra to bile acids, sometimes yielding high cosine similarity scores (>0.7), yet they lack key low-intensity diagnostic ions. Due to the low intensity of these ions, their presence or absence typically does not significantly affect spectral matching scores. However, our MassQL isomer filters can identify these MS/MS spectra. Therefore, we used the original bile acid MassQL queries^1^ to identify that 929 MS/MS spectra had the two characteristic fragment ions for mono-, di-, and trihydroxy bile acids or a spectral match to non-candidate bile acid spectral MS/MS reference libraries on GNPS2^32^. A portion of the 929 MS/MS spectra are adducts, in-source fragments, and multimers, overall reducing the number of unique bile acid structures. Based on retention time and peak shape correlation analysis in MZmine4^33,34^, a strategy that allows the discovery of ion forms associated with a specific molecule, we estimate these to represent 601 bile acids. Even though the data contained evidence for hundreds of unique bile acids, the stereo- and regiochemistry of hydroxylations are not defined.

The application of our MassQL filtering tree allowed us to further refine structural details within the bile acid core for a subset of the data. The first layer of the MassQL is the separation of mono-, di- and trihydroxylated bile acids and their ketones. A univariate analysis showed more unique monohydroxylated bile acids in feces and fewer di- and trihydroxy bile acids compared to the small intestinal fluids (**Supplementary Figure S3**). As we move down the MassQL filtering tree, we can now hypothesize the regio- and, in many cases, the stereochemistry of their hydroxyl positions. With the current implementation of the tree we can predict the bile acid cores of 178 spectra belonging to 111 unique bile acids. The highest number of spectral matches that could be assigned are dihydroxylated bile acids, symbolized here as (OH)_2_ (**Figure 2a**). These were categorized into individual bins of 3,12α-(OH)_2_, 3,6-(OH)_2_, 3,7-(OH)_2_, and 3- or 7-,12β-(OH)_2_ isomers following the application of the dihydroxy isomer filtering tree. Peak area distributions of each of these isomer groups varied across sample types, with the 3,7-(OH)_2_ isomer primarily consisting of the host-derived chenodeoxycholic acid steroid core, being more abundant in intestinal fluid. Similarly, the 3- or 7-,12β-(OH)_2_ isomers were also more prevalent in small intestinal fluid. Dihydroxy bile acids, such as 3,6-(OH)_2_, originally identified in pigs^35^, were found at higher levels in human feces than in intestinal samples. This is consistent with the recent work where hyodeoxycholic acid, 3α,6α-(OH)_2_ was measured to be around 700 nmol/g in pig feces compared to less than 1 nmol/g in the small intestine and 3 nmol/g in liver^36^. Microbially modified 3,12α-(OH)_2_ bile acids appeared in both intestinal fluid and feces, with taurine, glycine, serine, and aspartate conjugates being higher in the small intestine and ornithine and citrulline conjugates more abundant in feces consistent with observations in previous literature^37,38^.

**Figure 2.**
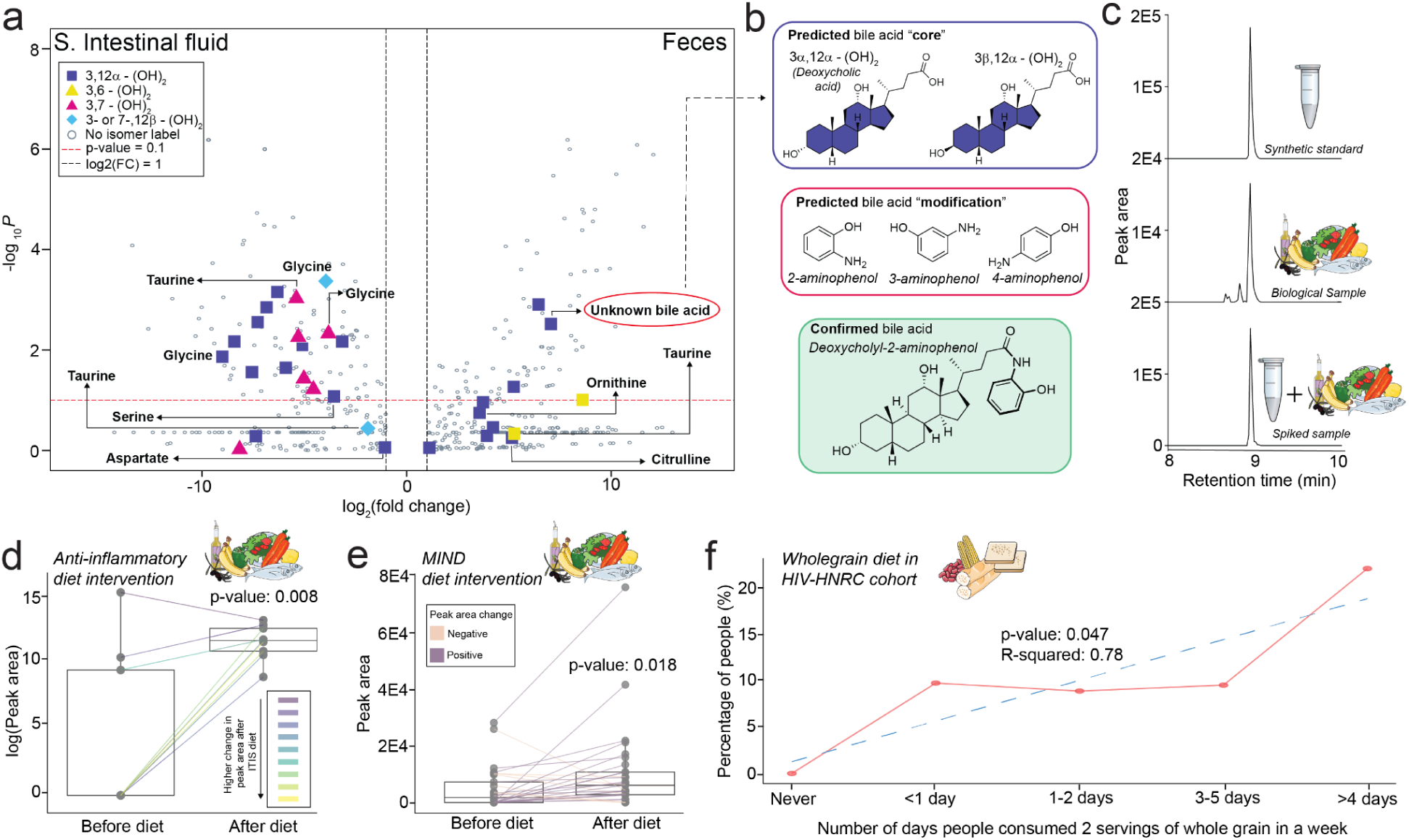
Discovery and Validation of Deoxycholyl-2-Aminophenol in Biological Samples. **a)** Distribution of bile acids (n=776; bile acids with more than 2-fold change from the total 929) between small intestinal fluid and fecal samples. **b)** Assignment of the bile acid steroid core by MassQL queries and putative identification of the modification using high-resolution MS1 data. **c)** Validation of deoxycholyl-2-aminophenol through retention time matching in biological samples using the synthetic standard. **d)** Peak area abundances of deoxycholyl-2-aminophenol before and after anti-inflammatory ITIS diet intervention. **e)** Peak area abundances of deoxycholyl-2-aminophenol before and after MIND diet intervention. **f)** Percentage of participants with non-zero peak area abundances of deoxycholyl-2-aminophenol. A non-parametric Wilcoxon signed-rank test was performed for the boxplots in Figures 2d and 2e. Horizontal lines indicate the median value, the first (lower) and third (upper) quartiles are represented by the box edges, and vertical lines (whiskers) indicate the error range, which is 1.5 times the interquartile range.

Within the 111 unique bile acid matches to this data where we could refine the stereo and regiochemistry of the core, there are 76 structurally uncharacterized bile acid modifications. For example, one of the 3,12α-(OH)_2_ bile acid isomers was elevated in feces compared to the intestine that also had the signature of a dihydroxylated bile acids at *m*/z*s* 321.26 and 339.27 and exhibited a mass shift of *m/z* 73.03 from the unmodified dihydroxylated bile acid^1^. The retention time analysis with other ion forms revealed this to be an in-source fragment of a dihydroxylated but modified bile acid. Considering the dehydration and proton gain, this mass change on the bile acid was 110.0606 Da. This ion is detected in the MS/MS spectrum (**Supplementary Figure S4a**) and corresponds to a molecular formula of C_6_H_7_NO. Searching for the modification’s high-resolution mass in the Human Metabolome Database (HMDB)^39^, and considering the structures that could be possible with the molecular formula, we narrowed down the possibilities that the modification attached to the bile acid core was most likely an aminophenol. Although only 3-aminophenol and 4-aminophenol were matched in HMBD, we cannot exclude the possibility of 2-aminophenol that was not present in HMDB. With only two possible bile acid cores - either 3α,12α- (OH)_2_ or 3β,12α-(OH)_2_ - the options for the final structure were narrowed to six candidates (**Figure 2b**). Although the lack of sample availability precluded retention time and ion mobility matching against synthetic standards in this dataset, we performed a MASST search^18^ to see if we could find other datasets containing this molecule for which samples were accessible. We found matches in 13 other fecal datasets, including three in-house studies for which the samples were available. Two datasets involved diet intervention studies. One was a pilot study on an anti-inflammatory ITIS diet (n = 17) to assess its effects on patients with active rheumatoid arthritis. The other was a larger study (n = 244) evaluating a combined Mediterranean and DASH diet (MIND) in cognitively normal older adults at risk for Alzheimer’s disease.We also obtained identical MS/MS spectral matches from the HIV Neurobehavioral Research Center (HNRC) cohort (n = 322) of people with and without HIV.

To confirm the predicted identity of the unknown bile acid, we synthesized the six possible bile acids and other similar structures, as controls, using multiplexed synthesis^9^. Using MS/MS, retention time and drift time information from ion mobility, we identified the unknown bile acid as 2-aminophenol conjugated to deoxycholic acid (3α,12α-(OH)_2_) in all three cohorts (**Supplementary Figures S4b, S4c**). We further validated the finding by synthesizing the pure standard of deoxycholyl-2-aminophenol as this allowed us to get NMR confirmation of the synthetic product, and performed retention time and co-migration analysis in the MIND diet study (**Figure 2c, Supplementary Figures S5, S6**). In this study, the 3β,12α-(OH)_2_ epimer was also conjugated with 2-aminophenol (**Supplementary Figure S4d**). No matches were obtained for the 3- or 4-aminophenol. 2-Aminophenol sulfate is an established marker for the consumption of whole grains, wheat, and corn, which are metabolized in part to 2-aminophenol by microbes^40–42^. Further exploration of the relationship between the deoxycholyl-2-aminophenol and dietary factors revealed an association with whole-grain consumption in all three studies. In the rheumatoid arthritis study, an anti-inflammatory diet including whole grains such as rye bread over 15 days increased the proportion of fecal deoxycholyl-2-aminophenol (**Figure 2d**). In the longitudinal MIND diet study, we observed similar increases after transitioning to the diet with recommended increase of whole grains (**Figure 2e**). From the HIV-HNRC dataset, we found an increase in the detection frequency of this bile amidate with self-reported whole-grain consumption, starting from the reported consumption of zero/week to more than two servings for 4 days/week of whole grains (**Figure 2f**). Thus, the production of this formerly unannotated bile acid is affected by dietary patterns.

In conclusion, this study introduces the concept of using a MassQL MS/MS fragmentation-based filtering tree approach to enable precise regio- and stereochemical differentiation of bile acid isomers, which were subsequently validated against reference standards. The method expands the ability to annotate bile acids and has led to the discovery of bile acids like deoxycholyl-putrescine and deoxycholyl-2-aminophenol, and its 3β isomer amidate, which had previously been detected but not structurally characterized. By making annotated spectra publicly available and enabling the queries, this approach enhances bile acid analysis, offering innovative tools for exploring their metabolic roles and biological significance, including retro-analysis of isomer variations in existing public LC-MS/MS data.

Although we have demonstrated the proof-of-concept works for the MassQL fragmentation tree to distinguish bile acid isomers, there are important limitations to consider. First, the current queries are not comprehensive for all potential bile acid isomers in human biology, as they depend on the availability of pure standards. However, our collection of 48 taurine-conjugated bile acid isomers from which we generated the current queries is among the largest available. Expanding access to additional standards, including those with more than three hydroxyl groups on the steroid core, will enable the use of machine learning to better classify bile acid isomer patterns as the training set grows. Second, some queries have higher false-negative rates due to the reliance on very low-intensity ions, some of which may not be consistently detected, especially in low-concentration samples. This can lead to missing fragment ions, limiting the MassQL tree’s ability to narrow down candidate isomers. To mitigate this, we recommend using higher sample concentrations and/or optimizing fragmentation energies to maximize the relevant fragment ion intensities. Despite these challenges, this method advances the ability to assign bile acid stereochemistry and regiochemistry directly from MS/MS spectra when the relevant ions are present. More importantly, the MassQL filtering tree can be retroactively applied to publicly available datasets, facilitating additional future large-scale data science efforts in bile acid analysis. Finally, we anticipate that the concept of the MassQL fragmentation tree could be applied to other classes of molecules where it is hard to distinguish isomers from existing MS/MS spectral matching algorithms.

## Methods

### MS/MS data acquisition of taurine-conjugated bile acids

We generated 1 mM stock solutions of our in-house collection of 48 taurine-conjugated bile acids in 100% MeOH. The standard samples were injected (5 μL) into a Vanquish ultra-high-performance liquid chromatography (UHPLC) system coupled to a Q Exactive quadrupole Orbitrap mass spectrometer (Thermo Fisher Scientific, Waltham, MA). A Kinetex polar C18 column (2.1 × 100mm, 2.6 μm particle size, 100 A pore size; Phenomenex, Torrance) was employed with a SecurityGuard C18 column (2.1 mm ID) at 40 °C column temperature. The mobile phases (0.5 mL/min) were 0.1% formic acid in both water (A) and ACN (B) with the following gradient: 0-0.5 min 5 % B, 0.5-1.1 min 5-25 % B, 1.1-7.5 min 25-40 % B, 7.5-8.5 min 40-99 % B, 8.5-10 min 99 % B, 10-12 min 5 % B. The mass spectrometer was operated in positive heated electrospray ionization with the following parameters: sheath gas flow, 53 AU; auxiliary gas flow, 14 AU; sweep gas flow, 3 AU; auxiliary gas temperature, 400 °C; spray voltage, 3.5 kV; inlet capillary temperature, 269 °C; S-lens level, 50 V. MS1 scan was performed at *m/z* 150-1500 with the following parameters: resolution, 35,000 at *m/z* 200; maximum ion injection time, 100 ms; automatic gain control (AGC) target, 1E6. Up to 5 MS/MS spectra per MS1 scan were recorded under the data-dependent mode (dd-MS^2^) with the following parameters: resolution, 17,500 at *m/z* 200; maximum ion injection time, 150 ms; AGC target, 5.0E5; MS/MS precursor isolation window, *m/z* 1; isolation offset, *m/z* 0.5; normalized collision energy (NCE) of 45 %; minimum AGC for MS/MS spectrum, 2.5E4; apex trigger, 2 to 5 s; dynamic precursor exclusion, 8 s. The data was deposited in GNPS/MassIVE and is publicly available at MSV000092003.

Before choosing NCE 45 for the generation of the MassQL queries, we ran the bile acid standards with different collision energies (NCE 10, 20, 30, 40, 45, 50, 60) (**Supplementary Figure S7**). The MS/MS ions and the ratio of their relative intensity were consistently observed in MS/MS spectra acquired with different NCEs as long as the MS/MS fragment ions were detected.

### Generation and validation of MassQL queries

The MS/MS spectral library generated using data acquired on a Thermo Q-Exactive with an NCE 45 was used to create the MassQL queries. This energy was selected as the optimal value to achieve rich MS/MS fragmentation within the *m/z* range of 100 to 300, which contains the critical MS/MS fragments for isomer annotation originating from the bile acid core. The MS/MS fragmentation spectra were manually inspected sequentially to identify diagnostic fragmentation ions that separated the bile acid isomers. For dihydroxylated bile acids, we first applied the MassQL query from our previous publication^1^ to detect these bile acids using ions at *m/z* 321.26 and *m/z* 339.27. The MS/MS spectra were then divided into two bins: (i) those containing ketones and (ii) those with one hydroxyl group on the side chain and one on the core. Remaining MS/MS spectra with two hydroxyl groups on the steroid core were further categorized into two additional bins: one containing 3- or 7,12α-(OH)_2_ isomers, and another with the remaining dihydroxy bile acid isomers. This separation was achieved using diagnostic ions at *m/z* 201.163 and *m/z* 211.147, with their intensity ratios calculated. Subsequent bins were refined further, distinguishing individual regioisomers and, in some cases, stereoisomers. A similar strategy was applied to generate the MassQL queries for monohydroxylated and trihydroxylated bile acids.

The MassQL queries generated were validated against the publicly available GNPS bile acid-specific library called BILELIB19 (https://gnps.ucsd.edu/ProteoSAFe/gnpslibrary.jsp?library=BILELIB19). The queries were developed by analyzing MS/MS data obtained from a Q-Exactive instrument. However, to validate the MassQL queries, we used an existing bile acid spectral library containing data acquired from both Q-Exactive and Q-TOF instruments as a control. Isomer labels in the BILELIB19 were assigned using a customized script submitted to GitHub (*see Code availability*). The MassQL queries for bile acid isomer assignment are developed for bile amidates and therefore we tested the specificity of the queries by subsetting the BILELIB19 for only bile amidates. The false discovery rates (FDR) were calculated for each MassQL query (*custom script in Code availability*) and are provided in **Supplementary Table S2.**

#### How to use the Bile acid isomer MassQL queries with LC-MS/MS dataset

To use the isomer MassQL queries on any untargeted LC-MS/MS dataset we have provided a custom Python script (see code availability section). Any MGF file containing the list of MS/MS spectra from a LC-MS/MS dataset is provided as an input to this Python script. The output is a .tsv file which provides the list of MS/MS scan numbers captured by each of the MassQL queries. Next, we need to apply the sequential binning of the MS/MS scans by traversing each branch of the fragmentation trees to obtain the final scans, which fall into one of the terminal bins. [**NOTE**: The sequential binning is critical to reduce false hits. Please do not skip this step]. This step can be done in R or Python script by obtaining the union of the scans from all the isomer groups in each branch of the filtering tree. As an example, for getting MS/MS scans of 3,12α-(OH)_2_ we need to find the common MS/MS scans obtained in “Dihydroxy”, “3,12α-(OH)_2_; 7,12α-(OH)_2_” and “3,12α-(OH)_2_” steps. We have used this approach in the analysis of the four LC-MS/MS datasets in this study (**Figures 2a, 2d-f**) and the script called “common_scans.R” can be found in the folder “BA_isomer_MassQL_filters” deposited to GitHub (*see code availability*). Alternatively, the MassQL bile acid isomer library called “GNPS-MASSQL-BILE-ACID-ISOMER” (*vide infra*) available on GNPS2 as a propagated library can be used while running molecular networking jobs. However, caution will be needed to manually investigate the spectral matches to ensure the bile acid diagnostic peaks are detected in the MS/MS spectra, and annotations will need to be validated by matching retention time with synthetic standards.

### Generation of publicly available MassQL bile acid isomer library

MS/MS Spectra from the extracted results of the Stage 2 MassQL queries conducted on GNPS/MassIVE repository in our previous publication^1^ (which we accessed from the “Extract Results” tab on the status page of the completed MassQL job) were used for isomer assignment. These Stage 2 MassQL queries considered data deposited in the public domain up to 08/19/2022. Bile acid isomer MassQL queries for the terminal bins in each of the monohydroxy, dihydroxy and trihydroxy fragmentation-based filtering trees were used to bin the MS/MS spectra from the repository searches conducted, followed by clustering the binned MS/MS spectra using MS-Cluster^43^. Clustering was performed to reduce the number of redundant MS/MS spectra belonging to the same bile acids. To cluster the spectra, we used the molecular networking workflow in the GNPS2 environment with precursor ion mass tolerance of 0.02 Da and MS/MS fragment ion mass tolerance of 0.05 Da. We set the minimum cosine score to 0.7 and minimum matched fragment peaks to 6. The job links are provided in **Supplementary Table S2**. We next selected the MS/MS spectra with the highest total ion current (TIC) from each group of the clustered spectra and created a second-generation MS/MS spectral reference library with the most plausible predicted isomers. The library is publicly available at <u>gnps.ucsd.edu/ProteoSAFe/gnpslibrary.jsp?library=GNPS-MASSQL-BILE-ACID-ISOMER</u> and the code to generate the library is deposited in GitHub (*see code availability*).

### Reanalysis of public data from GNPS/MassIVE

#### I. Small intestinal fluid and feces from healthy participants (MSV000094551)

This study received ethical approval from the Personal Protection Committee on 23 February 2022 and 9 March 2023, as well as from the French National Agency for the Safety of Medicines and Health Products (ANSM) on 2 June 2022 and 20 March 2023. It is formally registered under the identifier NCT05477069. Once the data have been evaluated, participants who request a summary of the study results will receive the information.

##### Sample preparation

Small intestinal and fecal samples from 15 healthy participants in the study were extracted using the protocol described previously^44^. Briefly, samples were homogenized by vortexing and sonicating for 10 min, followed by the addition of cold methanol (1:4 (v/v)) spiked with d-leucine for protein precipitation. Samples were incubated on ice for 30 min and then centrifuged for 15 min (15,000g, 4 °C). Supernatants were separated from pellets and evaporated under nitrogen. Dry extracts were resuspended with UHPLC solvent (80% water, 20% MeOH, 1% ACN, 0.1% formic acid).

##### LC-MS/MS data acquisition

Samples were injected (5 μL) on a Vanquish Flex ultra-high-performance liquid chromatography (UHPLC) system coupled to a Q Exactive Plus Orbitrap mass spectrometer (Thermo Fisher Scientific, Waltham, MA). The parameters for chromatographic separation and MS/MS data acquisition were previously described in detail^44^.

##### Feature-Based Molecular Networking and Data Analysis

The raw MS/MS spectra were converted to mzML files using MSconvert (ProteoWizard)^45^, followed by feature extraction using MZmine 4.2.0 with the following parameters: For mass detection, the factor of lowest signal was 4 for MS1 and 2.5 for MS2. For chromatogram building, the mass tolerance was set as 0.002 *m/z* or 10 ppm, the minimum consecutive scans as 4, and the minimum height as 5E4. Local minimum search for chromatographic deconvolution was performed with a minimum search range of 0.05 min, minimum ratio of peak top to side of 1.8, and maximum peak duration of 1.5 min. The peaks were de-isotoped within 5 ppm *m/z* and 0.05 minutes retention time tolerances, aligned with 0.0015 *m/z* or 5 ppm mass tolerance and 0.1 minutes retention time tolerance, then gap-filled with 5 ppm *m/z* tolerance and 0.1 minutes retention time tolerance. Ion forms were assigned using the metaCorrelate module with a retention time tolerance of 0.02 min, minimum feature height of 3E4 and intensity threshold for correlation at 1E4. Feature height correlation was selected using Pearson similarity measure with a minimum correlation of 70 % across a minimum of 3 samples.

The feature list was then exported as a feature quantification table (.csv) and an MGF spectra file. The feature quantification table was subsequently filtered with R scripts to remove features with average peak areas in samples lower than 3-fold of those in blanks. Metabolite annotations were performed with the default GNPS Spectral Libraries and GNPS-BILE-ACID-MODIFICATIONS library using the Feature-Based Molecular Networking (FBMN) workflow on GNPS2^46^. The spectra were filtered by removing MS/MS fragments within ±17 Da of the precursor m/z and to only keep the six most intense fragments in a ±50 Da window. The spectra were searched against the GNPS libraries with precursor and fragment ion mass tolerances of 0.02 Da and 0.05 Da, respectively, with a cosine score threshold of 0.6 and a minimum of 4 matched peaks. The GNPS2 job is available at: https://gnps2.org/status?task=4e5f76ebc4c6481aba4461356f20bc35.

The MGF output file from MZmine was subjected to distinct bins for bile acid isomers using the MassQL filters developed in this study using a custom Python script (*see code availability*). Therefore, a subset of the bile acid spectral matches from the FBMN job had regio- and stereochemistry defined for the bile acid steroid core. Using the “correlation group ID” which is an output of the ion identity molecular networking workflow in MZmine4, we also identified the potential adducts and in-source fragments. Next, univariate analysis was performed, and the results were plotted as a volcano plot showing the log2(FC) calculated for each bile acid in small intestinal fluid and feces. Fold change was calculated using MetaboanalystR 4.0 and plotted using the EnhancedVolcano package in R (https://github.com/kevinblighe/EnhancedVolcano). Only those bile acids with more than 2-fold change were shown on the volcano plot in **Figure 2a**. Amongst these, distinct shapes and colors highlighted the bile acids with regio- and stereochemistry assignment using the MassQL filters.

#### II. Anti-inflammatory diet intervention study in rheumatoid arthritis (MSV000084556)

A prospective, open-label pilot trial was conducted to evaluate the clinical and biological outcomes of a two-week isocaloric ITIS diet, with approval from the Institutional Review Board (IRB#161474).

##### Sample preparation

Extracts were prepared according to protocols published previously^47^ Briefly, 300 µL of MeOH-water (1:1) was added to each well using a multichannel pipette. The deep well plate was covered with a storage mat and sonicated for 5 minutes. The samples were placed in a 4°C refrigerator overnight to precipitate the proteins. Extracts were evaporated until dry using a CentriVap Benchtop Vacuum Concentrator (Labconco, Kansas City, MO, USA). The 96-well plates containing the dried extract were covered (96-deep well plate mats, Nunc 96 Well Caps for 1.0 mL Polystyrene DeepWell Plates) and stored at −80°C prior to analysis.

##### LC-MS/MS data acquisition

Fecal samples were analyzed using LC-MS/MS data acquisition which was performed on a Vanquish ultrahigh-performance liquid chromatography (UHPLC) system using a core-shell silica C18 column (2.1 x 50 mm, 1.7-μm particle size, 100-Å pore size; Kinetex, Phenomenex) coupled to a Q Exactive Orbitrap mass spectrometer (Thermo Fisher Scientific, Bremen, Germany). Five microliters of sample were injected and run at 0.5 ml/min on a gradient of solvent A (HPLC-grade water with 0.1% formic acid) and solvent B (HPLC-grade acetonitrile with 0.1% formic acid). The column was maintained at 40 °C. The UPLC elution gradient ran for 12.5 min per sample: 5% B from 0 min to 1 min, a linear gradient of 5 to 100% B over 8 min, a hold at 100% B for 2 min, a return to 5% B over 0.5 min, and a hold at 5% B for 2 min to equilibrate the column for the next sample. The flow was directed into a heated electrospray ionization source operated in positive ionization mode with the following parameters: an auxiliary gas flow rate of 14 arbitrary units (a.u.), sweep gas flow rate of 3 a.u., sheath gas flow rate of 52 a.u., spray voltage of +3.5 kV, capillary temperature of 270°C, auxiliary gas heater temperature of 435°C, and S-Lens RF level of 50. The data-dependent acquisition mode was used to acquire the data in which MS1 scans from *m/z* 100 to 1,500 (scan rate, 7 Hz) were followed by an MS2 scan, specifically a product ion scan produced using stepped normalized collision energy higher-energy collisional dissociation, of the five most abundant ions from the prior MS1 scan.

##### Feature-Based Molecular Networking and Data Analysis

The raw MS/MS spectra were converted to mzML files using MSconvert (ProteoWizard), followed by feature extraction using MZmine 4.2.0 with the same parameters as mentioned for Small intestinal fluid and feces from healthy participants (MSV000094551). Metabolite annotations were performed with the default GNPS Spectral Libraries and GNPS-BILE-ACID-MODIFICATIONS library using the Feature-Based Molecular Networking (FBMN) workflow on GNPS2. The spectra were filtered by removing MS/MS fragments within ±17 Da of the precursor *m/z* and to only keep the six most intense fragments in ±50 Da window. The spectra were searched against the GNPS libraries with precursor and fragment ion mass tolerances of 0.02 Da and 0.05 Da, respectively, with a cosine score threshold of 0.6 and a minimum of 4 matched peaks. The GNPS2 job is available at: https://gnps2.org/status?task=1234bd8eb0f940feba6170da30a07e36.

The MGF output file from MZmine was subjected to distinct bins for bile acid isomers using the MassQL filters developed in this study using a custom Python script (*see code availability*) to obtain regio and stereochemistry assigned bile acid annotations. Boxplots depicting the change in the peak area of the bile acid, deoxycholyl-2-aminophenol, in the diet intervention study was plotted in **Figure 2d** using the ‘ggplot2’ package in R (v 4.4.1) with R script provided in the code availability section.

#### III. Mediterranean-DASH diet intervention study (MSV000088646)

The Institutional Review Board at Rush University Medical Center approved the study protocol before the initiation of the study (ORA#17042801), and all participants provided written informed consent.

##### Sample preparation

A portion of the thawed fecal samples from the wooden swabs was transferred to new Eppendorf tubes. The extracts were prepared by adding 1 mL of MeOH:H_2_O (1:1) with 1 μM of sulfamethazine as internal standard, and the samples were homogenized in a Qiagen TissueLyzer II for 5 min at 25 MHz. The samples were incubated at −20 °C for 30 min to precipitate the proteins and then centrifuged for 5 min (14,000 g). Without disturbing the pellet, 100 μL of the supernatant was transferred to a shallow well plate and dried with a vacuum centrifuge concentrator (room temperature; ∼5 hours) and stored at −80 °C. Before instrumental analysis, the plates were redissolved by 10-minute sonication in 200 μL of 50% acetonitrile (v/v) with 100 μg/L sulfadimethoxine as the internal standard. Plates were centrifuged at 2000 g for 10 minutes, and 150 μL supernatants were collected for instrumental analysis.

##### LC-MS/MS data acquisition

The fecal extracts were injected (5 μL) into a Vanquish ultra-high-performance liquid chromatography (UHPLC) system coupled to a Q Exactive quadrupole orbitrap mass spectrometer (Thermo Fisher Scientific, Waltham, MA). A Kinetex polar C18 column (150 × 2.1 mm, 2.6 μm particle size, 100 A pore size; Phenomenex, Torrance) was employed with a SecurityGuard C18 column (2.1 mm ID) at 30 °C column temperature. The mobile phases (0.5 mL/min) were 0.1% formic acid in both water (A) and ACN (B) with the following gradient: 0-1 min 5 % B, 1-7 min 5-100 % B, 7-7.5 min 100 % B, 7.5-8 min 99-5% B, 8-10 min 5% B. The mass spectrometer was operated in positive heated electrospray ionization with the following parameters: sheath gas flow, 53 AU; auxiliary gas flow, 14 AU; sweep gas flow, 3 AU; auxiliary gas temperature, 400 °C; spray voltage, 3.5 kV; inlet capillary temperature, 269 °C; S-lens level, 50 V. MS1 scan was performed at *m/z* 100-1500 with the following parameters: resolution, 35,000 at *m/z* 200; maximum ion injection time, 100 ms; automatic gain control (AGC) target, 5.0E5. Up to 5 MS/MS spectra per MS1 scan were recorded under the data-dependent mode with the following parameters: resolution, 35,000 at *m/z* 200; maximum ion injection time, 100 ms; AGC target, 5.0E5; MS/MS precursor isolation window, *m/z* 3; isolation offset, *m/z* 0.5; normalized collision energy, a stepwise increase from 20 to 30 to 40 %; minimum AGC for MS/MS spectrum, 5.0E3; apex trigger, 2 to 15 s; dynamic precursor exclusion, 10 s.

##### Feature-Based Molecular Networking and Data Analysis

The raw MS/MS spectra were converted to mzML files using MSconvert (ProteoWizard). Feature detection and extraction was performed via MZmine 3.9 via batch processing. Data was imported using MS1 and MS2 detection with lowest noise signal set to 3 and 1.1 respectively. Sequentially, mass detection was performed and only ions acquired between 0.5 and 8 min, with MS1 and MS2 noise levels set to 5E4 and 1E3 respectively. Chromatogram builder parameters were set at 5 minimum consecutive scans, 1E5 minimum absolute height, and 10 ppm for *m/z* tolerance. Smoothing was applied before local minimum resolver, which had the following parameters: chromatographic threshold 85%, minimum search range retention time 0.2 min, minimum ratio of peak top/edge 1.7. Then, 13C isotope filter and isotope finder were applied. Features were aligned using join aligner with weight for *m/z* set to 3 and retention time tolerance set to 0.2 min. Features not detected in at least 3 samples were removed before performing peak finder. The feature list was then exported as a feature quantification table (.csv) and an MGF spectra file. Metabolite annotations were performed with the default GNPS Spectral Libraries and GNPS-BILE-ACID-MODIFICATIONS library using the Feature-Based Molecular Networking (FBMN) workflow on GNPS2. The spectra were filtered by removing MS/MS fragments within ±17 Da of the precursor *m/z* and to only keep the six most intense fragments in a ±50 Da window. The spectra were searched against the GNPS libraries with precursor and fragment ion mass tolerances of 0.02 Da and 0.05 Da, respectively, with a cosine score threshold of 0.6 and a minimum of 4 matched peaks. The GNPS2 job is available at: https://gnps2.org/status?task=8bfb999be1ee4747bb78d9c7015e34b5.

The MGF output file from MZmine was subjected to distinct bins for bile acid isomers using the MassQL filters developed in this study using a custom Python script (*see code availability*) to obtain regio and stereochemistry assigned bile acid annotations. Boxplots depicting the change in the peak area of the bile acid, deoxycholyl-2-aminophenol, in the diet intervention study was plotted in **Figure 2e** using the ‘ggplot2 package in R (v 4.4.1) with R script provided in the code availability section.

#### IV. HIV Neurobehavioral Research Center (HNRC) study (MSV000092833)

The UCSD Human Research Protections Program (irb.ucsd.edu) approved all study procedures before the initiation of the study and all participants provided written informed consent. All procedures involving human participants adhered to the ethical standards established by the institutional and/or national research committee (UCSD Human Research Protections Program, HNRC IRB#172092).

##### Sample preparation

Fecal samples were prepared with an automatic pipeline for simultaneous metagenomics-metabolomics extractions^48^. For the metabolite extraction, swabs were added into Matrix Tubes (ThermoFisher Scientific, MA, USA) containing 400 μL of 95% (v/v) ethanol and capped with the automated instrument Capit-All (ThermoFisher Scientific, MA, USA). The tubes were shaken for 2 minutes (1,200 rpm, the SpexMiniG plate shaker) followed by centrifugation for 5 minutes (2,700 g). The supernatant was then transferred to a deep well plate using an 8-channel pipette, dried with a vacuum centrifuge concentrator (room temperature; ∼5 hours) and stored at −80 °C. Before instrumental analysis, the plates were redissolved by 10-minute sonication in 200 μL of 50% acetonitrile (v/v) with 100 μg/L sulfadimethoxine as the internal standard. Plates were centrifuged at 450 g for 10 minutes, and 150 μL supernatants were collected for instrumental analysis.

##### LC-MS/MS data acquisition

The fecal extracts were injected (5 μL) into a Vanquish ultra-high-performance liquid chromatography (UHPLC) system coupled to a Q Exactive quadrupole orbitrap mass spectrometer (Thermo Fisher Scientific, Waltham, MA). A Kinetex polar C18 column (150 × 2.1 mm, 2.6 μm particle size, 100 A pore size; Phenomenex, Torrance) was employed with a SecurityGuard C18 column (2.1 mm ID) at 30 °C column temperature. The mobile phases (0.5 mL/min) were 0.1% formic acid in both water (A) and ACN (B) with the following gradient: 0-1 min 5 % B, 1-7 min 5-99 % B, 7-8 min 99 % B, 8-8.5 min 99-5% B, 8-10 min 5% B. The mass spectrometer was operated in positive heated electrospray ionization with the following parameters: sheath gas flow, 53 AU; auxiliary gas flow, 14 AU; sweep gas flow, 3 AU; auxiliary gas temperature, 400 °C; spray voltage, 3.5 kV; inlet capillary temperature, 269 °C; S-lens level, 50 V. MS1 scan was performed at *m/z* 100-1500 with the following parameters: resolution, 35,000 at *m/z* 200; maximum ion injection time, 100 ms; automatic gain control (AGC) target, 5.0E4. Up to 5 MS/MS spectra per MS1 scan were recorded under the data-dependent mode with the following parameters: resolution, 17,500 at *m/z* 200; maximum ion injection time, 100 ms; AGC target, 5.0E4; MS/MS precursor isolation window, *m/z* 3; isolation offset, *m/z* 0.5; normalized collision energy, a stepwise increase from 20 to 30 to 40 %; minimum AGC for MS/MS spectrum, 5.0E3; apex trigger, 2 to 15 s; dynamic precursor exclusion, 10 s.

##### Feature-Based Molecular Networking and Data Analysis

The raw spectra were converted to mzML files using MSconvert (ProteoWizard) followed by feature extraction using MZmine 4.8.0 with the following parameters: For mass detection, the noise factor was 3.5 for MS1 and 2.5 for MS2. For chromatogram building, the mass tolerance was set as 0.01 m/z or 10 ppm, the minimum consecutive scans as 3, and the minimum height as 5E2. Chromatograms were smoothed with the Savitzky Golay algorithm, followed by a local minimum search for chromatographic deconvolution with a minimum search range of 0.2 min, minimum ratio of peak top to side of 2, and maximum peak duration of 0.5 min. The peaks were de-isotoped within 5 ppm *m/z* and 0.05 minutes retention time tolerances, aligned with 0.01 *m/z* or 20 ppm mass tolerance and 0.3 minutes retention time tolerance, then gap-filled with 20 ppm *m/z* tolerance and 0.3 minutes retention time tolerance. The feature list was blank-subtracted with 300% fold change increases compared with maximum blank values and 30 minimum number of detection in blanks. The feature list was then exported as a feature quantification table (.csv) and an MGF spectra file and used without post-filtering. Metabolite annotations were performed with the default GNPS Spectral Libraries and GNPS-BILE-ACID-MODIFICATIONS library using the Feature-Based Molecular Networking (FBMN) workflow on GNPS2. The spectra were filtered by removing MS/MS fragments within ±17 Da of the precursor *m/z* and to only keep the six most intense fragments in ±50 Da window. The spectra were searched against the GNPS libraries with precursor and fragment ion mass tolerances of 0.02 Da and 0.05 Da, respectively, with a cosine score threshold of 0.6 and a minimum of 4 matched peaks. The GNPS2 job is available at: https://gnps2.org/status?task=c68fc90125424231bece7655426f9e17.

The MGF output file from MZmine was subjected to distinct bins for bile acid isomers using the MassQL filters developed in this study using a custom Python script (*see code availability*) to obtain regio and stereochemistry assigned bile acid annotations. The percentage of detection of the bile acid, deoxycholyl-2-aminophenol, in the study participants was plotted in **Figure 2f** using the ‘ggplot2 package in R (v 4.4.1) with R script provided in the code availability section. A linear model was fitted using the lm() function in R to calculate the p-value and coefficient of determination (R-squared).

### Multiplexed synthesis of deoxycholyl-putrescine

The synthetic standard for deoxycholyl-putrescine was obtained from the multiplexed reaction of deoxycholic acid with a mixture of amines^9^. Bile acid (deoxycholic acid; 0.08 mmol, 2.5 eq), and DMF (2.5mL) were added to a 20mL scintillation vial with a magnetic stir bar. To the solution solid EDC (0.2mmol, 5eq), neat DIPEA (0.3 mmol, 10 eq), and solid DMAP (0.2 mmol, 20 mg, 5 eq) were added sequentially and the solution was stirred at 23 °C. After 15 minutes, the corresponding amine compounds (mixture of 20 amino acids, 13 decarboxylated amino acids/polyamines including putrescine; 1 eq each) were added simultaneously and the reaction was stirred for 14hr. The mixture was concentrated in vacuo and used directly for LC-MS/MS analysis.

### Synthesis of bile acid conjugated with aminophenols

The synthetic standards of bile acid conjugated with 2-aminophenol, 3-aminophenol and 4-aminophenol were synthesized according to the following procedure. Each reaction consisted of one bile acid and one of the aminophenols. We used 5 different bile acid steroid cores for the reactions - deoxycholic acid 3α,12α-(OH)_2_; 3β,12α-(OH)_2_; chenodeoxycholic acid 3α,7α-(OH)_2_; ursodeoxycholic acid 3α,7β-(OH)_2_; hyodeoxycholic acid 3α,6α-(OH)_2_. Bile acid (0.08 mmol, 2.5 eq), and DMF (2.5 mL) were added to a 20 mL scintillation vial with a magnetic stir bar. To the solution solid EDC (0.2 mmol, 5 eq), neat DIPEA (0.3 mmol, 10 eq), and solid DMAP (0.2 mmol, 20 mg, 5 eq) were added sequentially and the solution was stirred at 23 °C. After 15 minutes corresponding amine compound (1 eq) were added simultaneously and the reaction was stirred for 14 h. The mixture was concentrated *in vacuo* and used directly for LC-MS/MS analysis.

### Synthesis and purification of deoxycholyl-2-aminophenol

Solid deoxycholic acid (0.5 mmol, 200 mg, 1 eq.) and 2 mL of DMF were added to a 20 mL scintillation vial with a magnetic stir bar. To this solution, solid HATU (0.6 mmol, 232 mg, 1.2 eq.) and neat DIPEA (0.7 mmol, 133 μL, 1.5 eq) were subsequently added, and the solution was stirred at 23°C. After 15 minutes, 2-aminophenol (0.5 mmol, 139 mg, 1 eq.) was added, and the reaction was stirred for 14 hours. The mixture was then concentrated in vacuo and purified by silica gel column chromatography. Elution with DCM:MeOH (8:2) afforded deoxycholyl-2-amino-phenol.

NMR spectra were collected at 298 K on a 600 MHz Bruker Avance III spectrometer fitted with a 1.7 mm triple resonance cryoprobe with z-axis gradients. (^1^H NMR: MeOD (3.31) at 600 MHz; ^13^C NMR: DMSO:CDCl_3_ (49.00) at 151 MHz). All shifts are reported in parts per million (ppm) referenced to the proton or carbon of the solvent. Coupling constants are reported in Hertz (Hz). Data for ^1^H-NMR are reported as follows: chemical shift (ppm, reference to protium; s = single, d = doublet, t = triplet, q = quartet, dd = doublet of doublets, m = multiplet, coupling constant (Hz), and integration).^1^H NMR (600 MHz, MeOD) δ 3.79 (s, 1H), 7.54 (d, 1H), 7.00 (t, 1H), 6.86 (d, 1H), 6.81 (t, 1H), 3.56 – 3.48 (m, 1H), 2.54 – 2.47 (m, 1H), 2.41 – 2.34 (m, 1H), 1.95– 1.85 (m, 6H), 1.85 – 1.74 (m, 3H), 1.67 - 1.56 (m, 4H), 1.57-1.35 (m, 9H), 1.38 – 1.23 (m, 5H), 1.23 – 1.05 (m, 4H), 1.01-0.89 (m, 4H), 0.75 (s, 3H). Data for ^13^C-NMR (DMSO:CDCl_3_): δ 172.59, 128.21, 126.18, 124.42, 121.73, 120.83, 118.74, 71.05, 69.91, 59.30, 48.70, 47.29, 46.05, 45.84, 43.98, 41.46, 35.45, 34.85, 33.55, 33.01, 32.83, 31.18, 27.96, 26.55, 25.61, 25.26, 25.12, 24.84, 23.33, 22.63, 16.35, 16.33, 15.85, 11.95.

### Co-migration analysis of deoxycholyl-putrescine

The reaction mixture with the standard of deoxycholyl-putrescine was dissolved in 100% MeOH and used to perform retention time and MS/MS spectral matching with the lion feces samples that were extracted previously for another study^1^. The fecal sample, synthetic standard, and sample spiked with the synthetic standard were subjected to LC-MS/MS analyses. The LC-MS/MS analyses were conducted with a Vanquish UHPLC system coupled to a Q-Exactive Orbitrap mass spectrometer (Thermo Fisher Scientific, Bremen, Germany). The chromatographic separation was performed on a Polar C18 column (Kinetex C18, 100 x 2.1 mm, 2.6 μm particle size, 100A pore size – Phenomenex, Torrance, USA), and the mobile phase consisted of H_2_O (solvent A), and ACN (solvent B), both acidified with 0.1% formic acid. The following gradient was employed to evaluate retention time matching between the synthetic standard and the compound present in the sample: 0-0.5 min 5% B, 0.5-1.1 min 5-30% B, 1.1-5.0 min 30-60% B, 5.0-9.0 min 60-100% followed by a 1.5 min washout phase at 100% B, and a 1.5 min re-equilibration phase at 5% B. The MS parameters are outlined above in the “MS/MS data acquisition of taurine-conjugated bile acids” section.

### Co-migration analysis of aminophenol bile amidates using LC-IM-MS/MS

Samples were resuspended in 200 µL of 1:1 MeOH/water with 1 µM of sulfadimethazine as internal standard. The chromatographic separation was performed by reversed-phase polar C18 (Kinetex Polar C18, 100 mm x 2.1 mm, 2,6 µm, 100 A pore size with a guard column, Phenomenex) using an Agilent UHPLC system coupled to a hybrid trapped ion mobility-quadrupole time-of-flight mass spectrometer (timsTOFpro2, Bruker Daltonics, Bremen, Germany). The mobile phase consisted of solvent A (water + 0.1% formic acid) and solvent B (ACN + 0.1% formic acid) and the column compartment was kept at 40 °C. Five microlitres of the samples were injected and eluted at a flow rate of 0.5 mL/min using the following gradient: 0 – 0.5 min 5% B, 0.5 – 1.1 min 25% B, 1.1 – 7.5 min 40% B, 7.5 – 8.5 min 99% B, 8.5 – 10 min 99% B, 10 – 10.1 min 5% B, 10.1 – 12 min 5% B. An Apolo II electrospray ionization source (ESI) was used to introduce ions into the TIMS analyzer. TIMS parameters were set as follows: 1RF potential of 300 Vpp, ion mobility was scanned from 0.45 to 1.45 Vs/cm^2^, accumulation and ramp time to 400 ms, ion charge control was set to 7.5E6. The TIMS dimension was pre-calibrated using four selected ions from the Agilent ESI LC/MS tuning mix [*m/z*, 1/K0: (322.0481, 0.7318 Vs cm−2), (622.0289, 0.9848 Vs cm−2), (922.0097, 1.1895 Vs cm−2), (1221,9906, 1.3820 Vs cm−2)] in positive mode. The QTOF was pre-calibrated using Na formate in positive mode at ppm standard deviation < 0.5%. Mass spectra were recorded at *m/z* 20–1300, and precursors for data-dependent acquisition were fragmented with an ion mobility-dependent collision energy, which was linearly increased from 30 to 50 eV in positive mode. The low-abundance precursor ions with an intensity above a threshold of 100 counts but below a target value of 4000 counts were repeatedly scheduled and otherwise dynamically excluded for 0.05 min. The data is publicly available on GNPS/MassIVE MSV000096291.

To validate collision cross-section and retention time, we extracted CCS and RT values for the synthesized standards and biological samples run in triplicates using Bruker Compass Data Analysis software. Using R (version 4.4.0), we made a scatter plot of the average CCS versus RT, as shown in **Supplementary Figure S3c**. Script is available on Github (*see Code availability*).

### Statistical analyses

The statistical analyses in the study were performed in R (version 4.4.1; R Core Team, 2024). For comparing the change in peak area abundances between the two timepoints in **Figures 2d and 2e** we performed a Wilcoxon signed-rank test. This non-parametric test was chosen due to the non-normal distribution of data. Statistical significance was determined using a two-sided test with a significance level of 0.05. Analyses were conducted in R using the wilcox.test function. For **Figure 2f**, linear models were fitted using the lm() function, and data manipulation was conducted with the dplyr package.

## Data availability

The annotated MS/MS spectra of bile acid isomers have been made publicly available under the name “GNPS-MASSQL-BILE-ACID-ISOMER” at <u>gnps.ucsd.edu/ProteoSAFe/gnpslibrary.jsp?library=GNPS-MASSQL-BILE-ACID-ISOMER</u>, providing a valuable resource for analyzing bile acid modifications and isomeric patterns. The library MGF is also archived at Zenodo (https://doi.org/10.5281/zenodo.14775114). Source data used for generation of the figures in the manuscript are available on GitHub (https://github.com/mohantyipsita/MassQL_bile_Acid_isomer_2025). The LC-MS/MS data acquired for the taurine conjugated bile acids used to develop the MassQL queries are available in GNPS/MassIVE with accession MSV000092003. A step-by-step guide to use the bile acid isomer spectral library is provided in the GNPS2 documentation (https://wang-bioinformatics-lab.github.io/GNPS2_Documentation/libraries/).

## Code availability

The Python code for isomer labels in BILELIB19, MassQL queries FDR validation and assigning the bile acid isomer labels to an input MGF file from any untargeted LC-MS/MS dataset is deposited and publicly available on GitHub (https://github.com/mohantyipsita/MassQL_bile_Acid_isomer_2025; https://github.com/Philipbear/BA_classification). All other R scripts used to generate the figures in the manuscript are available on GitHub (https://github.com/mohantyipsita/MassQL_bile_Acid_isomer_2025).

## Author contributions

Conceptualization, P.C.D.; methodology, P.C.D., I.M. and L.R.H.; formal analysis, I.M., S.X., V.C., J.A., Y.E.A., and P.C.D.; investigation, I.M., S.X., V.C., J.A., A.P., H.M.-R., S.Z., J.Z., A.T.; resources, A.L.G., T.S., C.X.W., J.E.L., M.S.A., R.J.E., D.J.M., D.R.F.J., M.S.C., M.G., R.M.V., R.K.D., D.S., M.W.; writing – original draft, P.C.D., I.M. and L.R.H.; writing – review & editing, all authors; supervision, P.C.D., funding acquisition, P.C.D., R.M.V.

## Acknowledgements

This project is supported by NIDDK 1R01DK136117-01 granted to P.C.D. The MIND data used in this project was enabled in part by the Alzheimer’s Gut Microbiome Project (AGMP), supported by the National Institute on Aging grants: 1U19AG063744 and 3U19AG063744-04S1, awarded to Dr. Kaddurah-Daouk at Duke University in partnership with multiple academic institutions. As such, the investigators within the AGMP not listed in this publication’s authors’ list, provided analysis-ready data, but did not participate in designing the study, conducting the analyses or writing of this manuscript. A listing of AGMP Investigators can be found at https://alzheimergut.org/meet-the-team/. A complete listing of the AD Metabolomics Consortium (ADMC) investigators can be found at: https://sites.duke.edu/adnimetab/team/. We also acknowledge funding from NIA R01AG056653 granted to R.M.V which collected samples from the MIND trial. M.G. and M.S.C were supported by a grant from the Krupp Endowed Fund. The HIV Neurobehavioral Research Center (HNRC) is supported by Center award P30MH062512 from NIMH to D.J.M. and R.J.E.

