## Supplemental Information for "MS/MS Mass Spectrometry Filtering Tree for Bile Acid Isomer Annotation"

**Supplementary Figure S1:** MS/MS fragmentation-based filtering tree for sequentially binning the regio- and stereoisomers of **a)** monohydroxy and **b)** trihydroxy bile acids with the structures shown for each step.

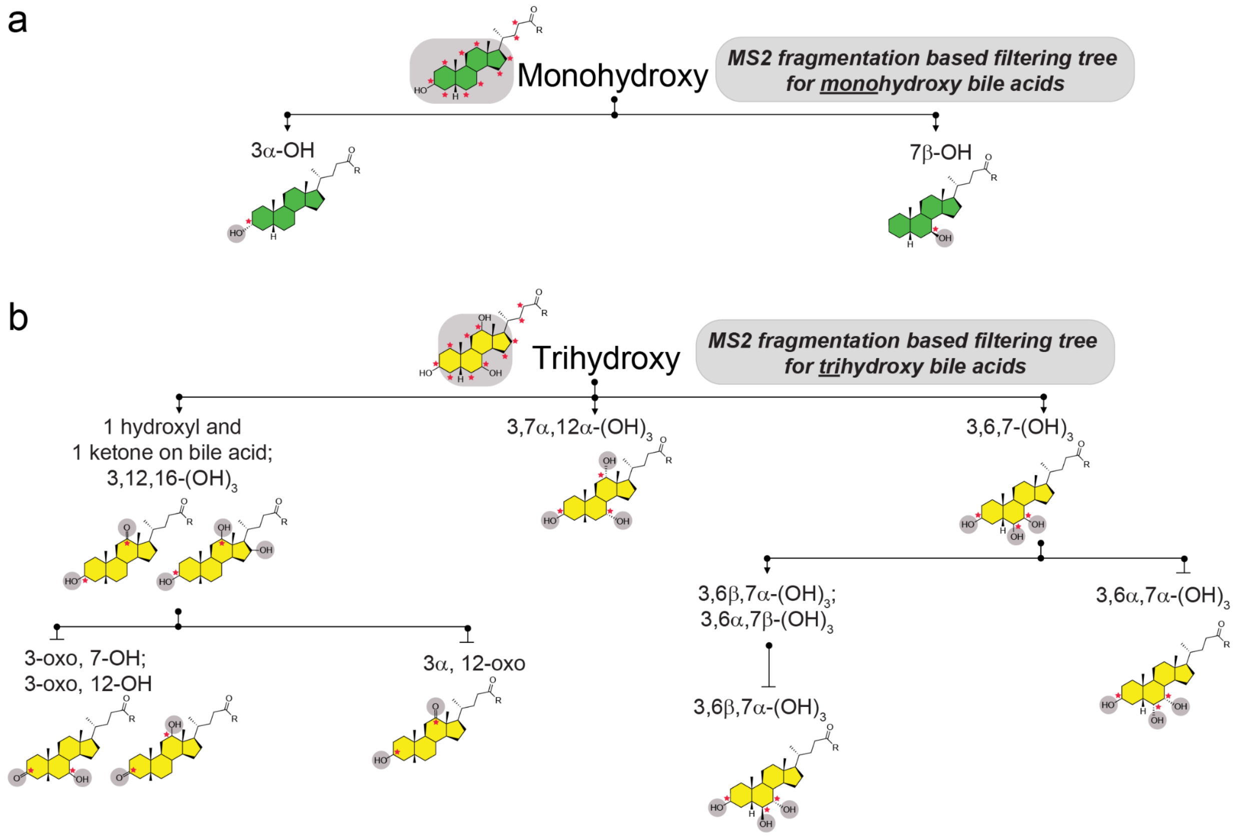

**Supplementary Figure S2:** Mirror plot of MS/MS fragmentation spectra of glycine conjugated keto bile acid (USI: mzspec:GNPS2:TASK-5d309e46ee8c4cfc835942dd1e10549d-nf_output/clustering/specs_ms.mgf:scan:843) and glycodeoxycholic acid (GDCA; USI: mzspec:GNPS:BILELIB19:accession:CCMSLIB00005435510) (Note: For the mono ketone bile acid, we used glycine conjugated mono ketone bile acid to develop the MassQL query due to lack of access to the taurine conjugated mono ketone bile acid).

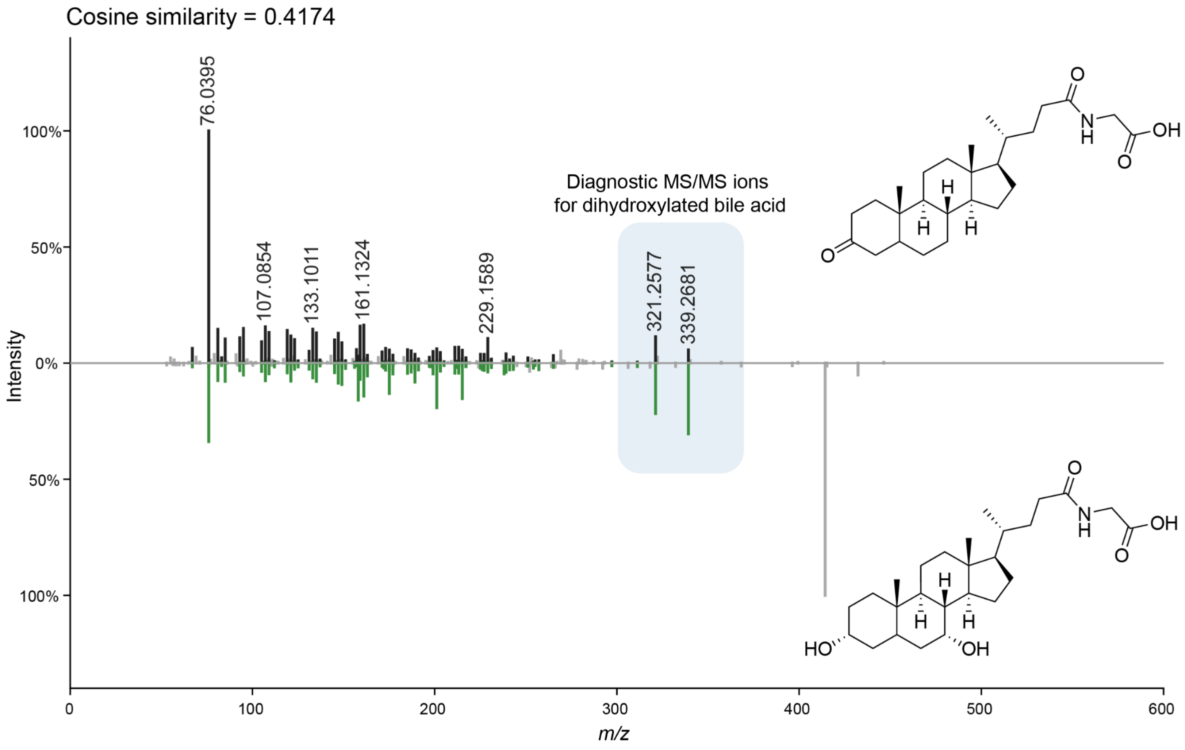

**Supplementary Figure S3:** Distribution of mono-, di-, and trihydroxy bile acids (n=776; bile acids with more than 2-fold change from the total 929) captured by the MassQL queries between small intestinal fluid and fecal samples.

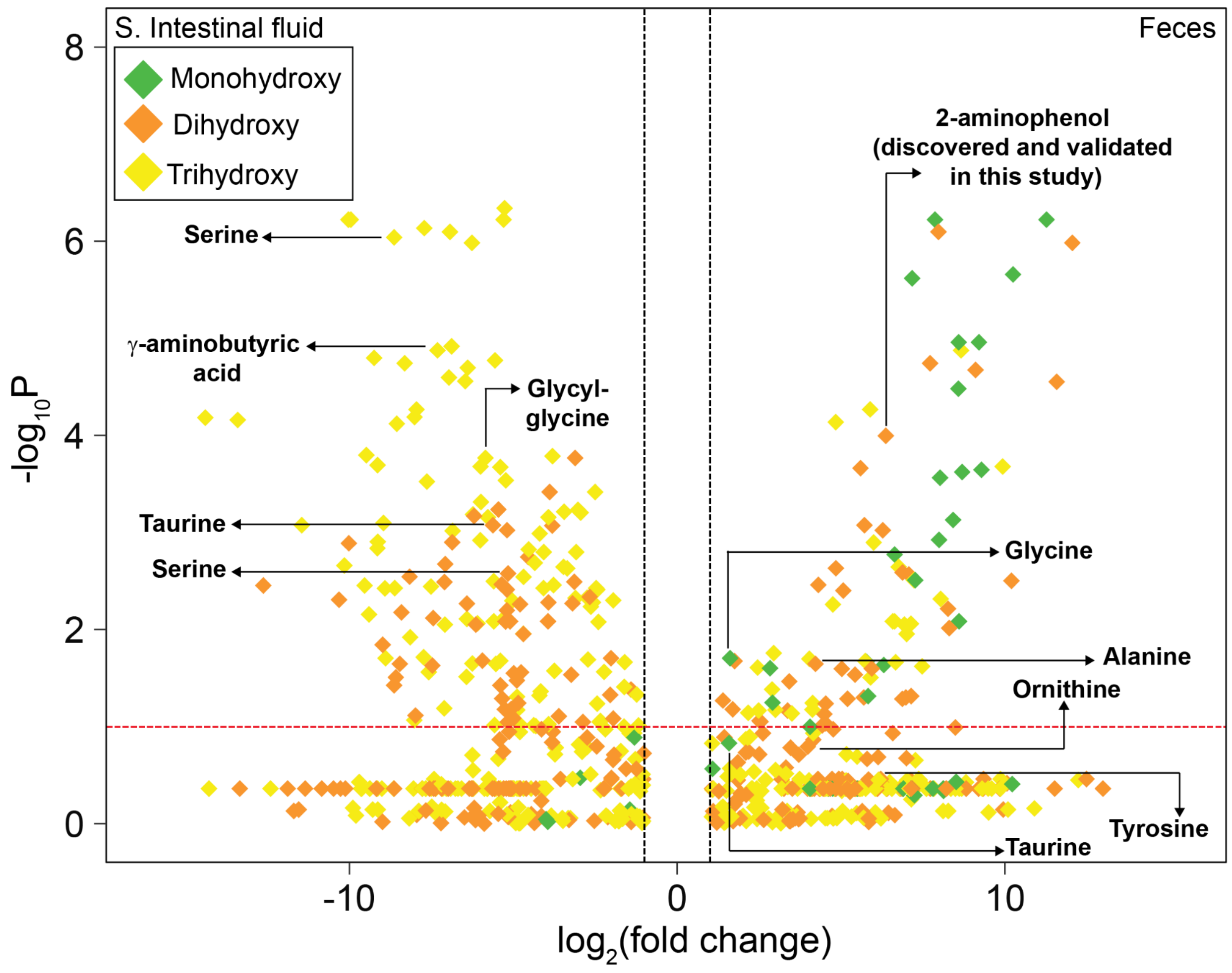

**Supplementary Figure S4: a)** MS/MS spectra of deoxycholyl-2-aminophenol (USI: mzspec:GNPS2:TASK-4e5f76ebc4c6481aba4461356f20bc35-nf_output/clustering/spectra_reformatted.mgf:scan:115873). **b)** Validation of deoxycholyl-2-aminophenol through ion mobilogram matching in biological samples using synthetic standard. **c)** Retention time and Collision Cross Section (CCS) values from ion mobility for the aminophenol conjugated bile acids in biological samples. **d)** Retention time match for 3β,12α-$(OH)_2$ conjugated with 2-aminophenol, the minor species.

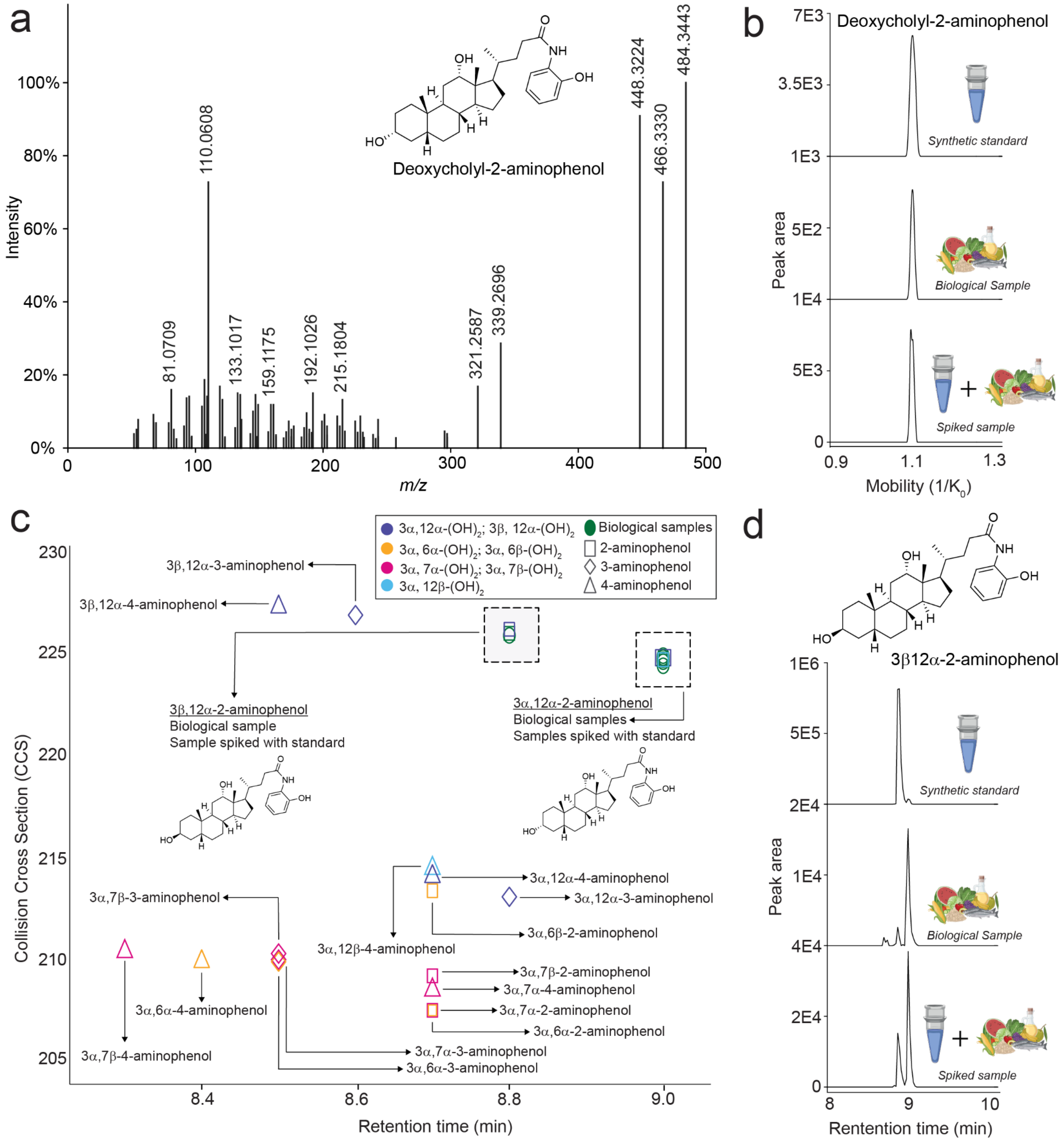

**Supplementary Figure S5.** $^{1}$H-NMR (600 MHz, MeOD) of the pure standard of deoxycholyl-2-aminophenol.

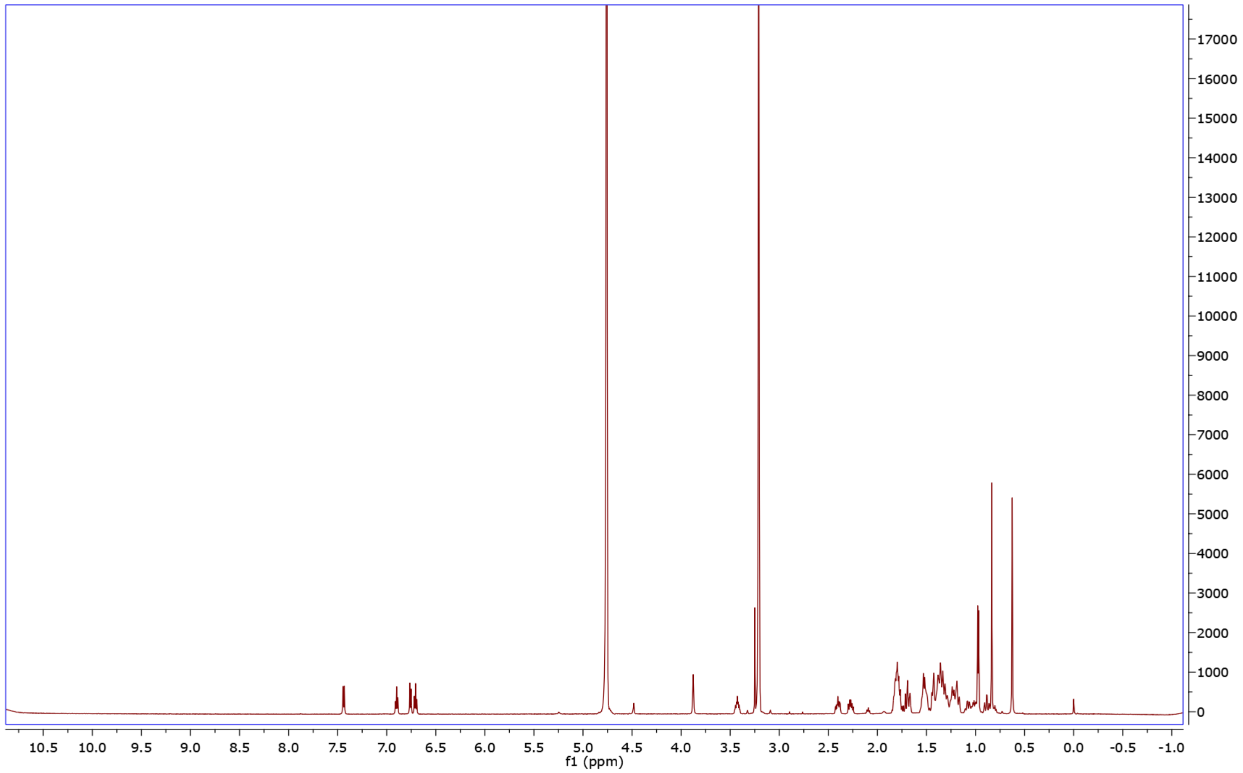

**Supplementary Figure S6.** $^{13}$C-NMR (151 MHz, DMSO:$CDCl_3$) of the pure standard of deoxycholyl-2-aminophenol.

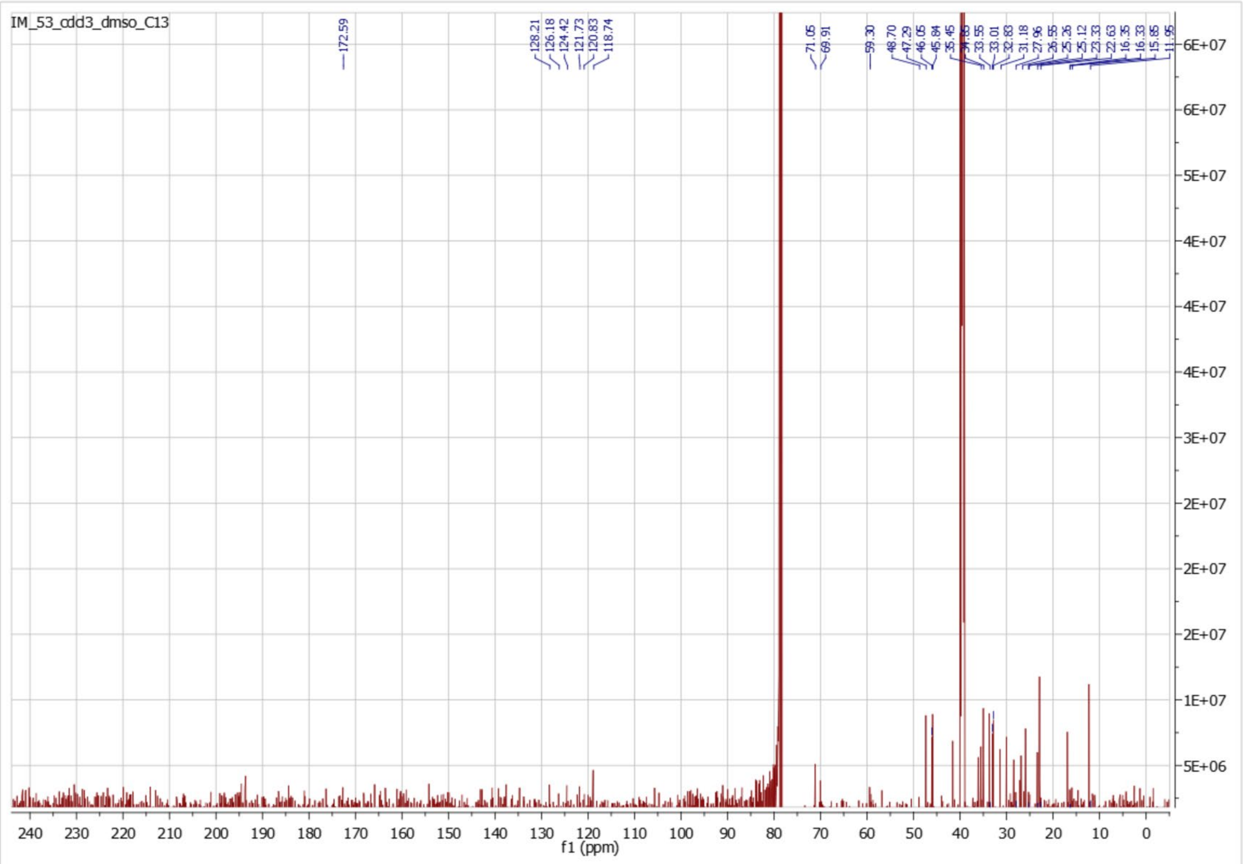

**Supplementary Figure S7.** MS/MS fragmentation ions used for calculating the intensity ratios for **a)** taurolithocholic acid, **b)** taurodeoxycholic acid, and **c)** taurocholic acid acquired on an Orbitrap at different fragmentation energies.

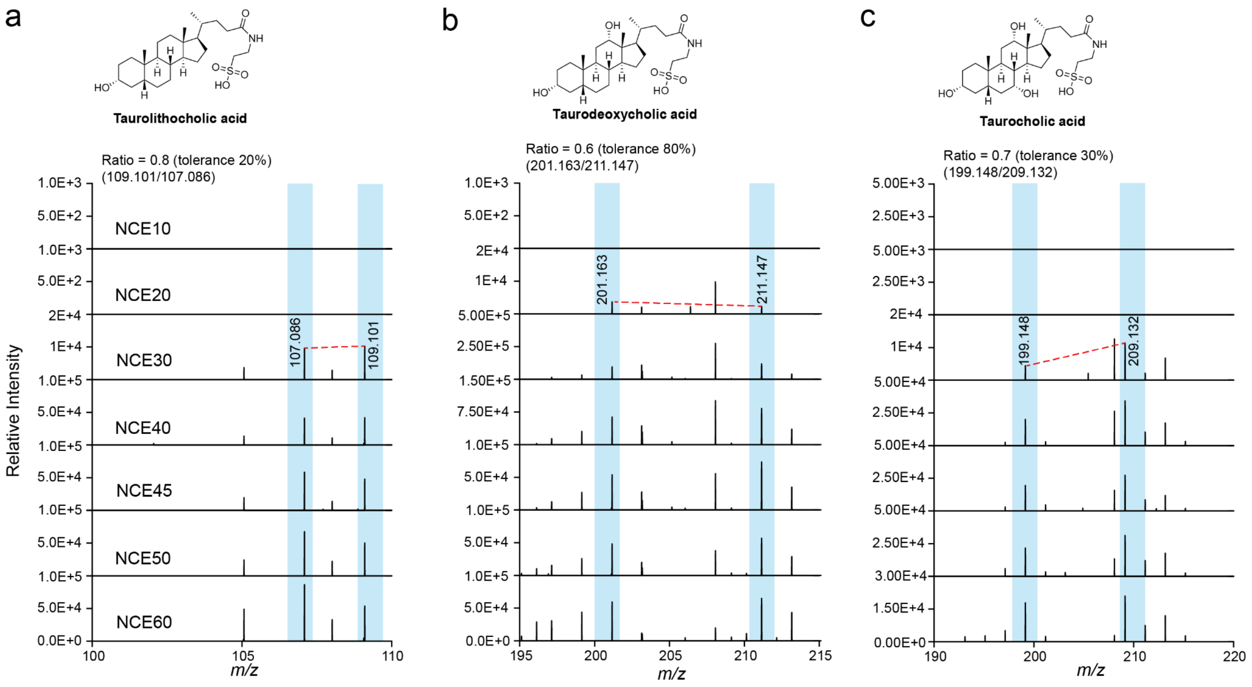
